# Effect of viewing distance on object responses in macaque areas 45B, F5a and F5p

**DOI:** 10.1101/2020.05.04.070862

**Authors:** I Caprara, P Janssen

**Author notes:** Corresponding Author: Peter Janssen; Irene Caprara (;). Caprara I - Division of Biology and Biological Engineering, Computation and Neural Systems, Caltech, Pasadena, CA 91125, USA.

## Abstract

To perform real-world tasks like grasping, the primate brain has to process visual object information so that the grip aperture can be adjusted before contact with the object is made. Previous studies have demonstrated that the posterior subsector of the Anterior Intraparietal area (pAIP) is connected to frontal area 45B, and the anterior subsector of AIP (aAIP) to F5a (Premereur et al., 2015). However, the role of area 45B and F5a in visually-guided object grasping is poorly understood. Here, we investigated the role of area 45B, F5a and F5p in visually-guided grasping. If a neuronal response to an object during passive fixation represents the activation of a motor command related to the preshaping of the hand, such neurons should prefer objects presented within reachable distance. Conversely, neurons encoding a pure visual representation of an object should be less affected by viewing distance. Contrary to our expectations, we found that the majority of neurons in area 45B were object- and viewing distance selective, with a clear preference for the near viewing distance. Area F5a showed much weaker object selectivity compared to 45B, with a similar preference for objects presented at the Near position emerging mainly in the late epoch. Finally, F5p neurons were less object selective and frequently preferred objects presented at the Far position. Therefore, contrary to our expectations, neurons in area 45B – but not F5p neurons – prefer objects presented in peripersonal space.

**Significance statement:** The current experiment provides the first evidence on the neural representation of distance in frontal areas that are active during visually-guided grasping. Area 45B and F5a neurons were object- and distance-selective, and preferred the near viewing distance even for objects with identical retinal size. In area F5p we observed strong visual responses with an unexpected preference for the Far viewing distance, suggesting that the motor-related object representation was still active during the presentation of objects outside reaching distance.

## Introduction

In the last decades, extensive research has been conducted to assess how different brain areas in the dorsal visual stream and its target areas in frontal cortex contribute to object grasping. Although neurons in parietal area V6A also respond during object grasping (Fattori et al., 2010; Fattori et al., 2012; Filippini et al., 2017), the most studied parieto-frontal network for controlling the hand comprises the Anterior Intraparietal Area (AIP) and the ventral premotor cortex (PMv). Neurons in AIP are selective for real-world objects (Murata et al., 2000), grip type (Baumann et al., 2009), 3D- (Srivastava et al., 2009) and 2D images of objects and very small fragments (measuring 1–1.5 deg – Romero et al., 2012, 2014). Overall, the target areas of AIP in frontal cortex seem to have similar properties. Neurons in the anterior subsector of PMv (F5a), which is effectively connected to the anterior subsector of AIP (aAIP – Premereur et al., 2015), respond selectively to images of 3D objects (Theys et al., 2012) and are active during object grasping. Visual-dominant neurons (i.e. responding during object fixation but not during grasping in the dark) are present in F5a (Theys et al., 2012) but not in F5p (Murata et al., 2000; Raos et al., 2006). A subset of neurons in the posterior part of PMv (F5p) are selective for real-world objects, even during passive fixation (Raos et al., 2006). In area F5p, objects are represented mainly in terms of the grip type used to grasp the object (Murata et al., 2000). Instead, area 45B, located in the anterior bank of the lower ramus of the arcuate sulcus and receiving input from the posterior subsector of AIP (pAIP – Premereur et al., 2015), responds selectively to 2D images of objects, with a preference for very small contour fragments, as in pAIP (Caprara et al., 2018).

While object selectivity has been extensively studied in parietal and premotor areas, few studies have investigated the neural representation of space at the single-cell level in parietal (Hadjidimitrakis et al., 2014, 2015) and frontal cortex (Bonini et al., 2014; Lanzilotto et al., 2016). Lesion studies in monkeys (Rizzolatti et al., 1983) and humans (Halligan and Marshall, 1991) have indicated the existence of distinct networks processing near and far space. In a recent fMRI study, Clery et al. (2018) have shown that visual stimuli appearing in near space activate temporal, parietal, prefrontal and premotor areas, whereas stimuli in far space produced activations in a different network spanning occipital, temporal, parietal, cingulate and orbitofrontal cortex. Although with some overlap, near and far space processing seemed to be segregated in two networks, as already suggested in human studies (Weiss et al., 2000; Aimola et al., 2012). To date, no study has investigated the other two subsectors of area F5 (F5a, F5p) and area 45B concerning space processing.

We wanted to investigate the encoding of viewing distance in three frontal areas receiving input from AIP, and to explore the nature of the object responses in these areas using objects presented at different distances with identical retinal size. If a neuron’s response to objects mainly reflects the motor plan to grasp that object (as in F5p), this response should be strongly modulated by viewing distance, since objects appearing in extrapersonal space should not activate the motor plan to the same degree. On the other hand, if a neuron encodes object information in visual terms (as we expect in 45B and in a subset of neurons in F5a), the neuronal response to objects in near and far space should be similar, provided that retinal size is constant.

## 3. Materials and methods

### 3.1 Subjects and Surgery

Two adult male rhesus monkeys (D, 7 kg and Y, 12.5 kg) served as subjects for the experiments. All experimental procedures, surgical techniques, and veterinary care were performed in accordance with the NIH Guide for Care and Use of Laboratory Animals and in accordance with the European Directive 2010/63/EU and were approved by the local ethical committee on animal experiments of the KU Leuven.

An MRI-compatible head fixation post and recording chamber were implanted during propofol anesthesia using dental acrylic and ceramic screws above the right arcuate sulcus in Monkey D and over the left arcuate sulcus in Monkey Y. The recording chamber allowed us to access 45B, F5a and F5p, as shown on MR images with a microelectrode in one of the recording positions for each area (Figure 1B).

**Figure 1.**
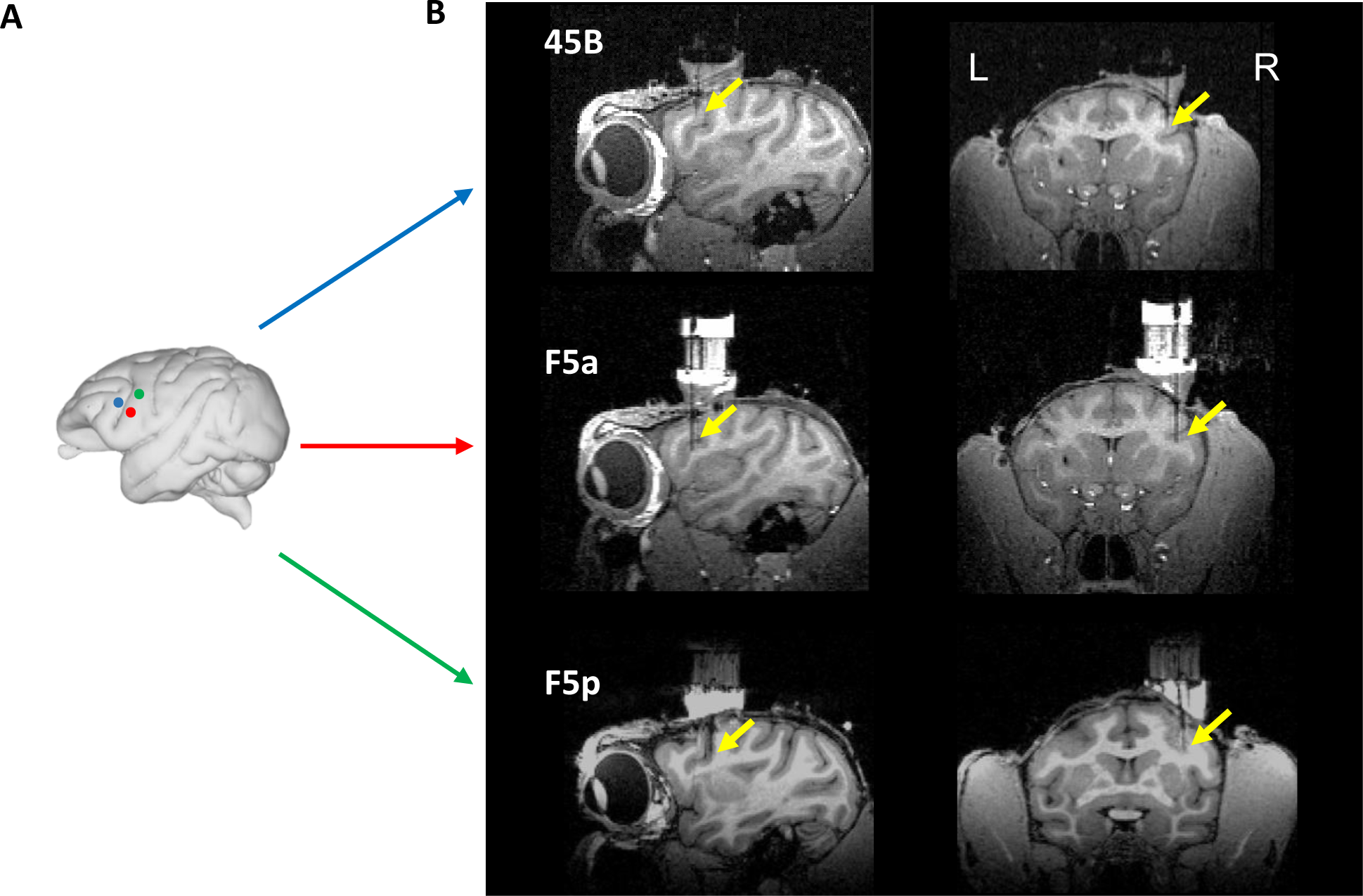
Anatomical location of areas of interest. A. Schematic view of a macaque brain (edited from the ‘Scalable Brain Atlas’ http://link.springer.com/content/pdf/10.1007/s12021-014-9258-x; Calabrese et al 2015). Colored dots represent the areas of interest in the arcuate sulcus, respectively Area 45B – blue, Area F5a – red – and Area F5p - green. B. Anatomical electrode recording position in Monkey D. Yellow arrows indicate the electrode tip location in the three areas. Equivalent recording position in Monkey Y were reported (data not shown).

### 3.2 Apparatus and Recording procedures

During the experiments, the monkey was seated upright in a chair with the head fixed. Each animal was trained not to move the other arm during the whole duration of the session. In front of the monkey, an industrial robot (Universal Robots, model UR-6-85-5-A) picked up the to-be-grasped object from a placeholder and presented it to the monkey. Four different objects (small plate 2 × 1 cm, small sphere 3 cm diameter, large plate 6 × 4 cm, large sphere 6 cm diameter) were pseudo-randomly presented one at a time in front of the monkey. The object could appear either at a Near position (28 cm viewing distance, at chest level ∼ 20 cm reaching distance measured from the center of the hand rest position to the center of the objects – peripersonal space) or at a Far position (57 cm – extrapersonal space – Figure 2). The average object luminances were: large sphere 3.3 cd/m2; small sphere 11.2 cd/m2; large plate and small plate 3.4 cd/m2. Since both object size and viewing distance were exactly two times larger at the Far position compared to the Near position, the retinal size of a small object at the Near position and the same-shaped large object at the Far position was identical.

**Figure 2.**
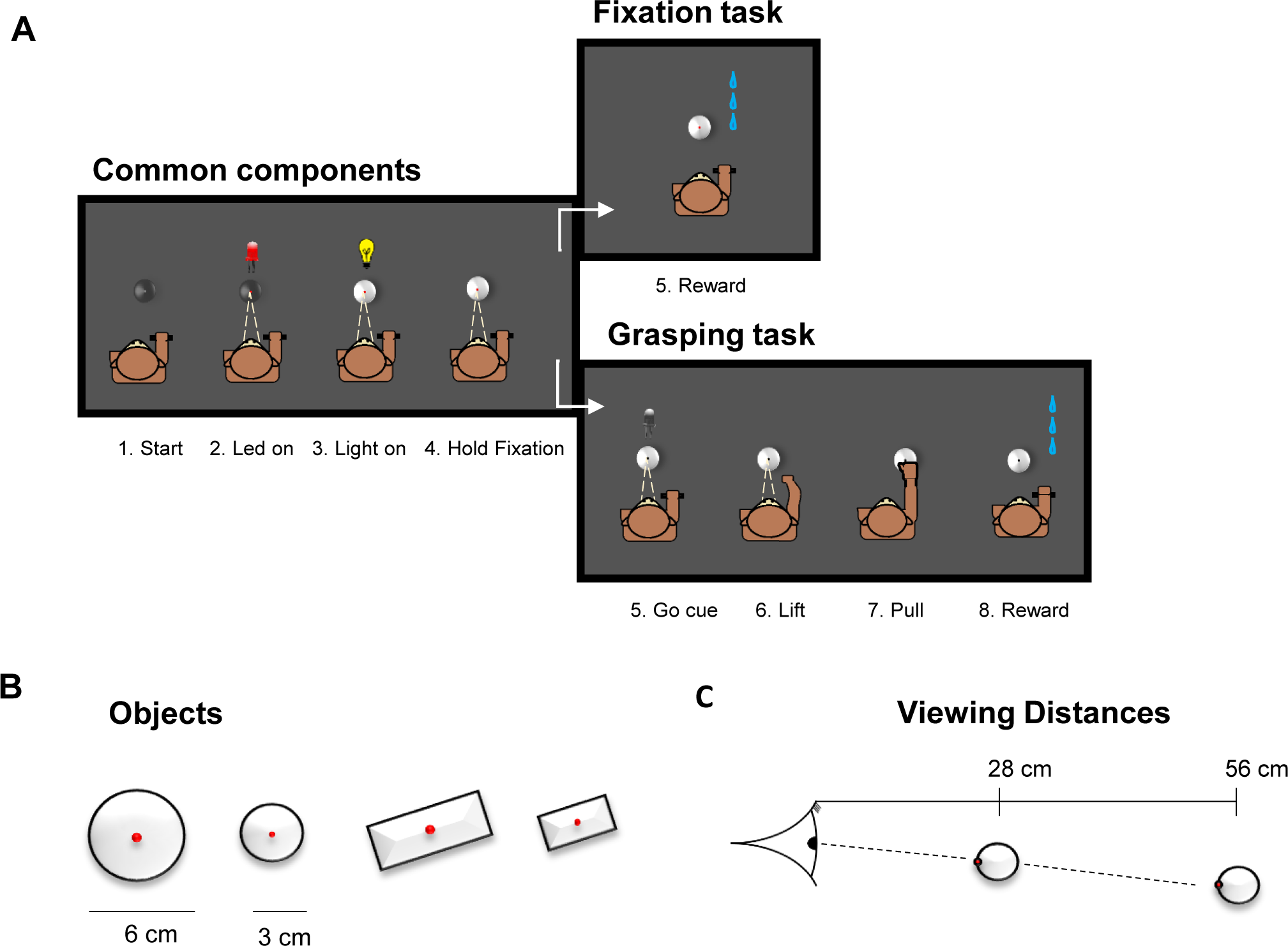
Schematic representation of the behavioral task, stimuli and distances. A. Behavioral task. The first block represents the common events to both Fix and Grasp tasks. The right-upper block corresponds to Fix task, the right-lower represents the Grasp task. B. Objects. large and small spheres and plates. C. Two viewing distances (Near, Far) from the monkey’s eyes. See details in the text.

The objects required different types of grasping depending on their size, but comparable across monkeys: a pad-to-side grip (for the small sphere and the small plate), and a finger-splayed wrap, corresponding to a whole-hand grip (for the large sphere and the large plate – (Macfarlane and Graziano, 2009). Fiber-optic cables detected the resting position of the hand, the start of the reach to grasp movement, and the pulling of the object. The start of the hand movement was detected as soon as the palm of the hand was 0.3 cm above the resting plane, whereas pulling of the object was detected when the object was pulled for 0.5 cm in the horizontal axis.

We recorded single-unit activity with standard tungsten microelectrodes (impedance, 1 MΩ at 1 kHz; FHC) inserted through the dura by means of a 23-gauge stainless-steel guide tube and a hydraulic microdrive (FHC). Neural activity was amplified and filtered between 300 and 5000 Hz. Spike discrimination was performed online using a dual time-window discriminator, and displayed with LabView and custom-built software. Spiking activity and photodiode pulses (corresponding to the onset of light in the object) were sampled at 20 kHz on a DSP (C6000 series; Texas Instruments, Dallas, TX). We continuously monitored the position of the left eye with an infrared-based camera system (Eye Link II, SR Research), sampling the pupil position at 250 Hz.

### 3.3 Behavioral Tasks

The two monkeys were trained to perform two tasks in a dark room, a passive fixation (Fix trials) and a visually guided grasping (VGG, Grasp trials) task (Figure 2). In Fix trials, the monkey had to passively fixate a small LED on the object which appeared either at the Near or at the Far distance until he received a juice reward. In Grasp trials, instead, he had to first fixate the LED on the object presented at the Near distance, and then, after a visual go cue (the offset of the LED on the object), to lift the hand from the resting position and pull the object in order to get the reward. Both types of tasks were performed using a robot, which picked one object at the time from a wooden placeholder, and presented it in front of the monkeys at one of the two distances.

To start both the Fix and Grasp trials, the monkey had to place the hand contralateral to the recording chamber in the resting position in complete darkness. During this time the robot picked an object from the box and moved it either to the Near or to the Far position. A red fixation LED inserted in the middle of the object was then illuminated, which the monkey had to fixate (keeping the gaze within a ±3.5 degrees throughout the trial). After 500 ms, a white LED illuminated the object from within for a variable amount of time (350 -1500 ms). If the red fixation LED did not dim until the end of the trial (Fix trials), the monkey’s gaze remained inside the electronically defined window, and the hand remained in the resting position, the animal received a juice reward. In the other half of the trials, the red LED in the middle of the object dimmed (Go cue – Grasp trials), which was the signal for the monkey to lift its hand from the resting position, reach and pull the object for 300 ms (holding time) in order to obtain a reward. Note that when the object appeared at the Near position, the animal could not predict whether the trial would be a Fix trial or a Grasp trial up to the moment of the dimming of the red fixation LED.

As a control, we recorded the activity of a small subset of neurons in one monkey during memory guided grasping (MGG). Similar to the VGG task, the monkey had to place its contralateral hand in the resting position in complete darkness to start the trial. During this time the robot picked an object from the box and present it at the Near position. A red fixation LED inserted in the middle of the object was then illuminated, which the monkey had to fixate (keeping the gaze inside a ±3.5 degree fixation window throughout the trial). After 500 ms, a white LED illuminated the object from within for 400 ms. After this fixed amount of time, the white LED dimmed, but the red LED stayed on. After a variable time (350-1500 ms), the red LED dimmed serving as a signal for the monkey to lift its hand from the resting position, reach and pull the object in total darkness for 300 ms (holding time) in order to obtain reward. Note that after the dimming of the red LED, no other sources of light were present in the room.

### 3.4 Data Analysis

All data analysis was performed in Matlab (Mathworks). For each trial, the baseline firing rate was calculated from the mean activity recorded in the 300 ms interval preceding Light onset. For the Fix trials, we then calculated the net neural responses by subtracting the baseline activity from the mean activity observed between 40 and 600 ms after Light onset. We tested visual responsivity by means of t-tests (p<0.05) comparing the baseline activity to the activity in the period after Light onset.

For the Fix conditions, both cell-by-cell and population analysis were performed to quantify distance and object selectivity. For every responsive neuron, we computed a two-way ANOVA with factors *[distance]* and *[object]*, and counted the number of cells with a significant main effect of distance, a significant main effect of object, or a significant *[distance x object interaction]*.

We plotted the averaged population net response to each object at the Near and at the Far distance, for each area. All the following statistical analyses were performed in two visual epochs, i.e. Early – from 40 to 200 ms after Light onset – and Late – from 200 to 400 ms.

To quantify object selectivity, we ranked the average net responses to the objects based on the odd trials at each distance (Near or Far) and separately for each area (test for significance in the two epochs – t test p<0.05). Then, we plotted the average responses in the even trials based on this ranking.

To test whether the object selectivity was similar at the two positions, we ranked the objects based on the responses at the Near distance and plotted the average responses to the same objects at the Far distance.

To assess distance selectivity, we first compared the average net response at the preferred distance (Best, i.e. the object eliciting the highest response in the test) to the average net response to the same object at the other distance (Worst). As in the previous analyses, we determined the preferred distance based on the even trials and plotted the responses in the odd trials. Then, to assess the preference of each area for a particular viewing distance, we compared the average net response to the preferred object presented at the Near position to the average net response to the same object at the Far position (test for significance in the two epochs – t test p<0.05). The same analysis was also repeated selecting the preferred object at the Far position. Finally, to assess the neural selectivity for objects with identical retinal size, we compared the average net response to the best small object presented at the Near position to the average net response to the same shaped large object at the Far distance (test for significance in the two epochs – t test p<0.05).

For Grasp trials in the light (VGG), we plotted the average response to the best object across all neurons per area, aligning the neural activity to multiple events during the trial (Light onset – Go cue – Lift – Pull). We measured the responsivity (t test p<0.05) in each epoch of the trial by comparing the baseline firing rate to the activity in each epoch after the above-mentioned events. We also analyzed the activity of a subset of neurons in each of the three areas during grasping in the dark (MGG).

## 4. Results

### Object and distance selectivity

All data included in this analysis were recorded in Fix trials, which were randomly interleaved with Grasp (VGG) trials. Since the results were highly similar between the two animals, we combined the data sets for all analyses. All neurons included (Monkey D: 45B, n = 57; F5a, n = 45; F5p, n = 44; Monkey Y: 45B, n = 57; F5a n = 44; F5p n = 33 – total number of neurons: 280) responded significantly to at least one object during fixation after Light onset. In each of the areas, we observed both Object- and Distance-selective neurons. The example neurons in Figure 3 illustrate the typical object (upper panel) and distance (lower panel) selectivity we observed in each of the areas. The example neuron of area 45B was clearly object-selective, and responded significantly stronger to the two large objects than to the two small objects (ANOVA, p = 0.02) at the Near distance. In F5a, the object selectivity was generally weaker and the responses evolved more slowly, as in the example neuron (middle column), whereas F5p neurons often showed transient responses to light onset with some object selectivity (right column). Note that in all three example neurons, the object selectivity was only apparent at the near position. The lower panel of Figure 3 illustrates examples of distance selectivity in the three areas without clear object selectivity. The example neuron of 45B preferred the Near position whereas the example neurons of F5a and F5p preferred the Far position.

**Figure 3.**
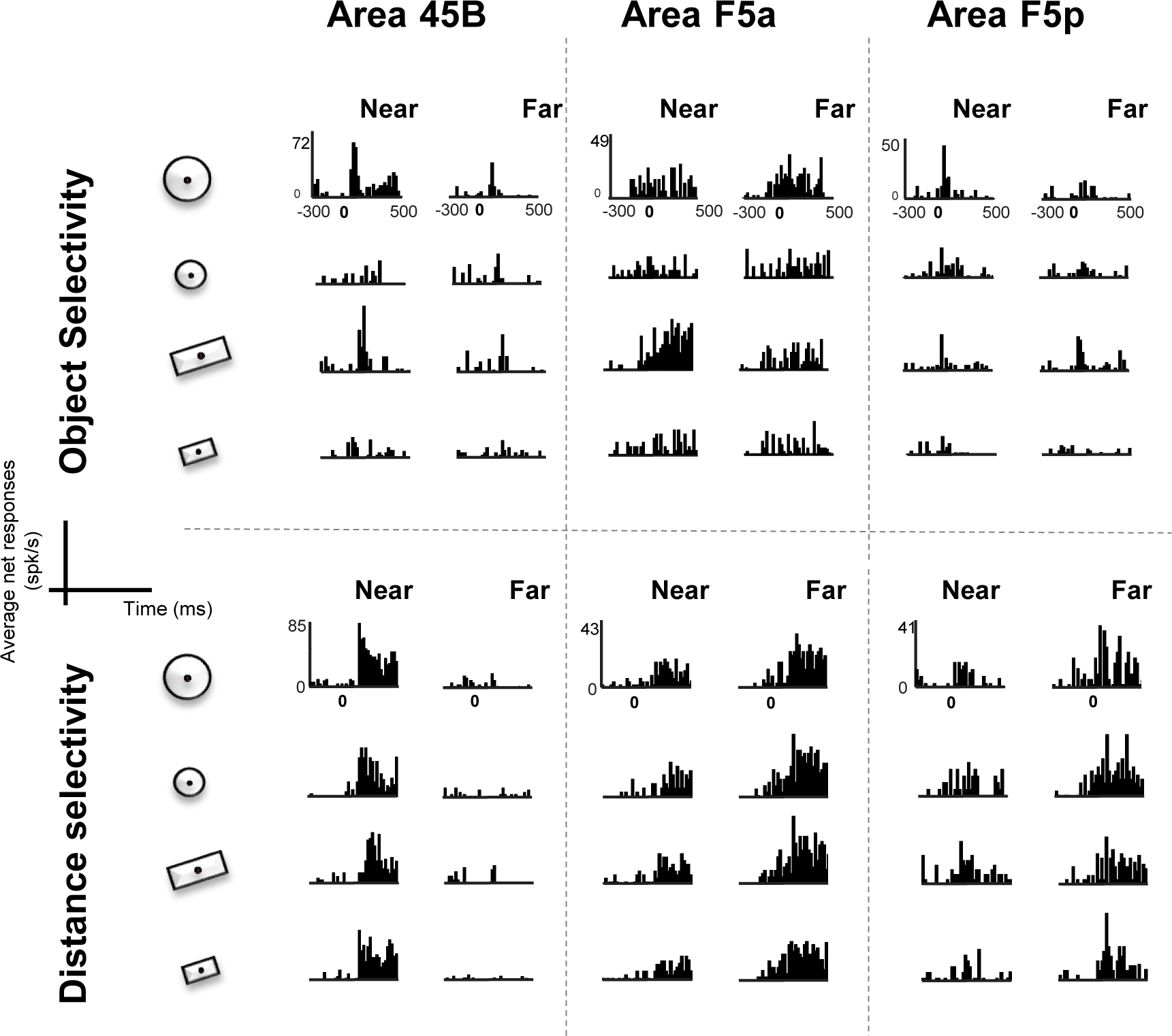
Example neurons in area 45B, F5a and F5p. In the upper panel, three neurons showing Object selectivity (from left to right, preference for large sphere, large plate and large sphere); in the lower panel, three examples of Distance selective neurons (from left to right, preference for Near, Far, Far).

The example neurons in Figure 3 also illustrate the different response profiles in the three areas. While area 45B neurons had a fast increase of the firing rate after object onset, neurons in area F5a showed a slower ramping up of the response without any brisk increase after object onset. Neurons in area F5p, instead, had an intermediate profile between 45B and F5a. To quantify the effects of Object, Distance and the interaction between these two factors, we performed a two-way ANOVA on the responses of each neuron. We observed a significantly higher proportion of object-selective neurons in area 45B compared to F5a and F5p (main effect of Object significant in 41% of the neurons in 45B, compared to 20% in F5a, p = 7.4 × 10^−4^; and 23% in F5p, p = 5.4 × 10^−3^, z-test). The proportion of neurons with a significant effect of Distance did not differ between the areas (45B: 44%; F5a: 36% and F5p: 38%, Table 1). Interestingly, of all selective neurons (significant at least for either *Object, Distance* or the *Object x Distance interaction*), a subset of neurons also had an effect of Size, preferring either the two large objects or the two small objects (45B: 18/74, 24%; F5a: 7/44, 16%; F5p: 5/42, 12%).

**Table 1.** Numbers of neurons with a significant effect of Shape, Distance and Interaction in 45B (N = 114), F5a (N = 89) and F5p (N = 77) – Two-way ANOVA.

|  | 45B | F5a | F5p |
| --- | --- | --- | --- |
| **Object** | 47 (41%) | 18 (20%) | 18 (23%) |
| **Distance** | 50 (44%) | 32 (36%) | 29 (38%) |
| **Interaction** | 23 (20%) | 12 (13%) | 9 (12%) |

To illustrate the degree of object selectivity in each area, we ranked the objects for each selective neuron (one-way ANOVA, separately at the Near and at the Far positions) based on the responses in the odd trials, and plotted the average response to the four objects in the even trials based on this ranking (Figure 4). In area 45B, we observed significant differences between preferred and non-preferred objects in the Early epoch (0-200ms) at both distances (ANOVA, main effect of object p = 3.05 × 10^−8^ and p = 3.53×10^−4^ Near and Far, respectively). In the Late epoch (200-400ms), instead, area 45B preserved a strong object selectivity at the Near distance but not at the Far (p = 2.17 × 10^−9^ and p = 0.03, respectively). Conversely, we did not observe comparable significant differences across objects in the other two areas (at the Near distance F5a was only significant during the late epoch p = 3,5 × 10^−3^, while F5p in the early epoch – p = 3,1 × 10^−3^; at the Far distance, p>0.05 at all-time epochs). A two-way ANOVA with factors [*object]* and *[area]* revealed that the object selectivity was significantly stronger in 45B than in the other two frontal areas, both at the Near (p = 2.19 × 10^−5^) and at the Far (p = 8.71 × 10^−6^) viewing distance.

**Figure 4.**
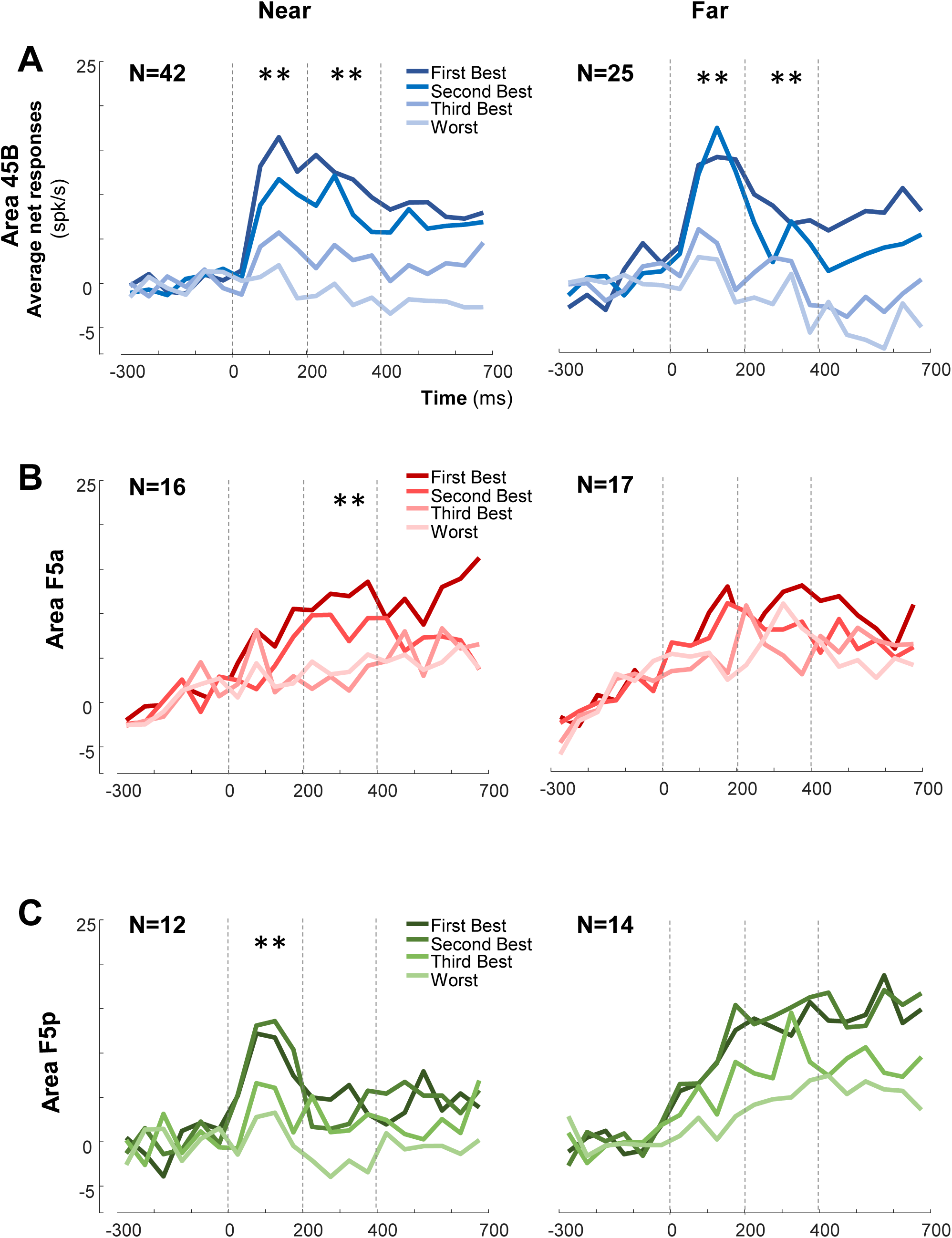
Object selectivity. Average ranked (on odd trials) population response of the even trials of object selective neurons at Near and Far distance (one-way ANOVA). Average net response across monkeys in area 45B (blue shades; N = 42 and N = 25 for Near and Far, respectively), F5a (red shades; N = 16 and N = 17 for Near and Far, respectively) and F5p (green shades; N = 12 and N = 14 for Near and Far, respectively). Independently from the color, the darkest shades represent the First Best object; progressively lighter colors represent lower ranking. The bin size was 50 ms. One asterisk indicate p<0.05; two asterisks indicate p<0.01.

Next, we investigated the effect of viewing distance. We first determined the preferred viewing distance for every neuron. Contrary to our expectations, we observed significantly more neurons preferring the near distance in area 45B (58%) and in area F5a (55%) than in F5p (39%, p = 5.1 × 10^−3^ and p = 0.02, z-test; Table 2). Thus, the majority of F5p neurons preferred the Far distance. Then, to quantify the strength of the distance selectivity, we compared the average net response to the preferred object at the Best distance to the response to that same object at the Worst distance (Figure 5). Area 45B and F5a had a significant effect of distance, both in the early and in the late epoch (p = 1.98 × 10^−7^ and p = 3.62 × 10^−9^, respectively early and late epoch in 45B; p = 4.49 × 10^−4^ and p = 2.30 × 10^−4^, respectively early and late epoch in F5a) while area F5p only showed a significant effect of distance in the late epoch (p = 2.24 × 10^−4^). Averaged across neurons and across the entire stimulus presentation interval, frontal neurons responded 79% (45B), 55% (F5a) and 74% (F5p) less to the preferred object at the worst distance compared to the same object at the best viewing distance.

**Table 2.** Preference for the two distances in area 45B, F5a and F5p

|  | 45B |  | F5a |  | F5p |  |
| --- | --- | --- | --- | --- | --- | --- |
|  | Near | Far | Near | Far | Near | Far |
| Large | 42 | 23 | 24 | 18 | 14 | 25 |
| Small | 24 | 25 | 25 | 22 | 16 | 22 |
| Total | 66 (58%) | 48 (42%) | 49 (55%) | 40 (45%) | 30 (39%) | 47 (61%) |

**Figure 5.**
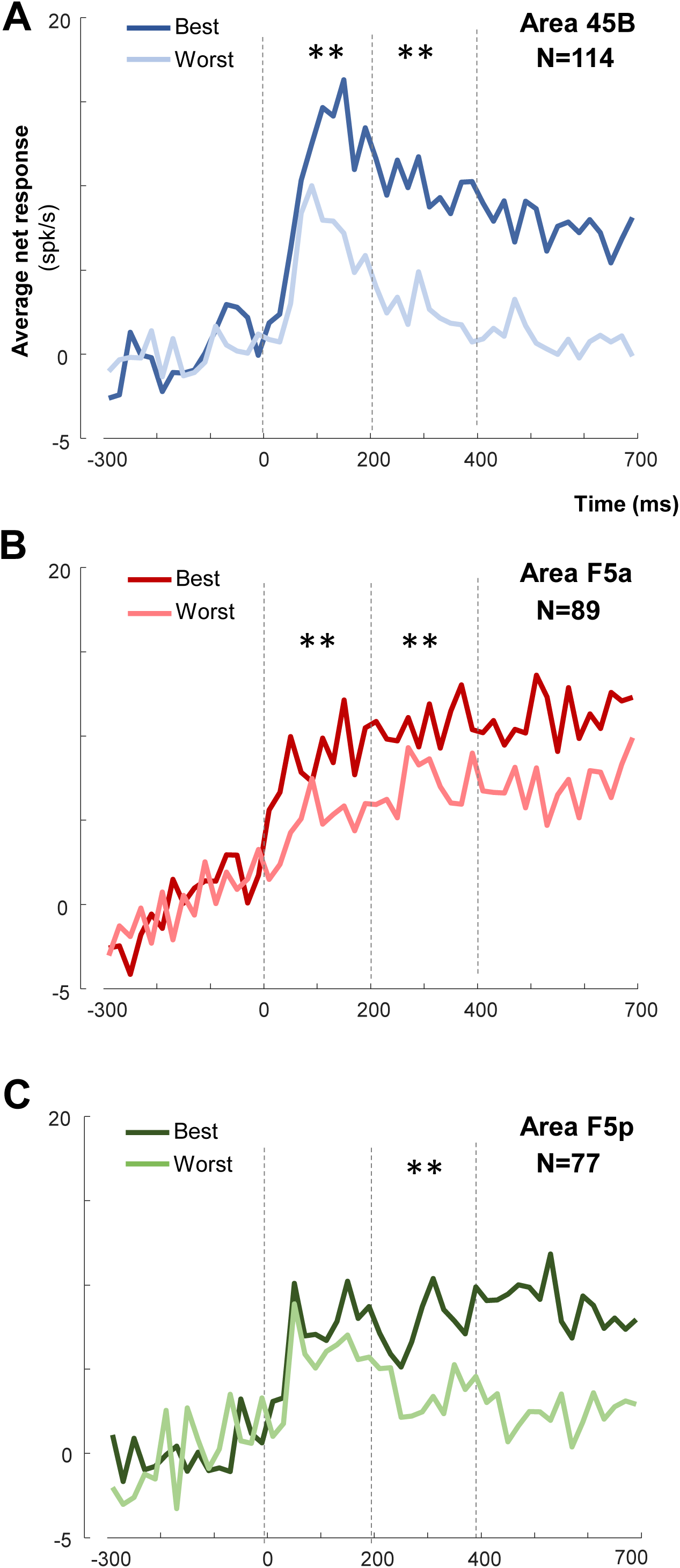
Distance selectivity comparing object responses at the Best and at the Worst distances. Average distance selectivity of the even trials, previously selected on odd trials, comparing the same object at the two distances (bin size = 20 ms): darker colors represent the Best distance for the three areas (blue for 45B, red for F5a, and green for F5p); the lighter color shade indicates Worst distance. Asterisks indicate statistical significance (p<0.01).

Viewing distance may not only affect the responses to the preferred object, but also the object preference of the neuron. In other words, is the object selectivity invariant across viewing distances? In order to quantify the invariance of the object selectivity across distance, we first ranked the objects based on the average response of each neuron at the Near position, and then calculated the response to the same objects at the Far position. When averaged across the population, the neuronal tuning for objects at the Near distance was stronger in 45B than in the other two areas: the slope of the regression line for area 45B was -3.7, for area F5a the slope was -2.6, and for F5p the slope was -2.4 spikes/sec/stimulus rank (Figure 6 and Table 3). However, in every area the object preference was only weakly preserved at the Far distance. The slopes of the regression lines at the Far position were -0.8 (CI: -1.4 – -0.2) in 45B, -0.4 (CI: -1 – +0.2) in F5a, and -0.3 (CI: -1.3 – +0.7) in F5p. Thus, neurons in these three areas are sensitive to changes in the viewing distance of an object, as relatively small alterations in object position (less than 30 cm) can produce very significant changes in selectivity.

**Table 3.** Slopes (spikes/sec/stimulus rank) and confidence intervals of the regression lines of the average Near ranked and on the Far on Near ranking responses, divided per area (45B, N =114; F5a, N = 89; F5p, N = 77).

|  | **45B** | **F5a** | **F5p** |
| --- | --- | --- | --- |
| **Near ranked** | -3.7 (-4.1 – -3.3) | -2.6 (-3.0 – -2.3) | -2.4 (-2.7 – -2.2) |
| **Far on Near ranking** | -0.8 (-1.4 – -0.2) | -0.4 (-1.0 – 0.2) | -0.3 (-1.3 – 0.7) |

**Figure 6.**
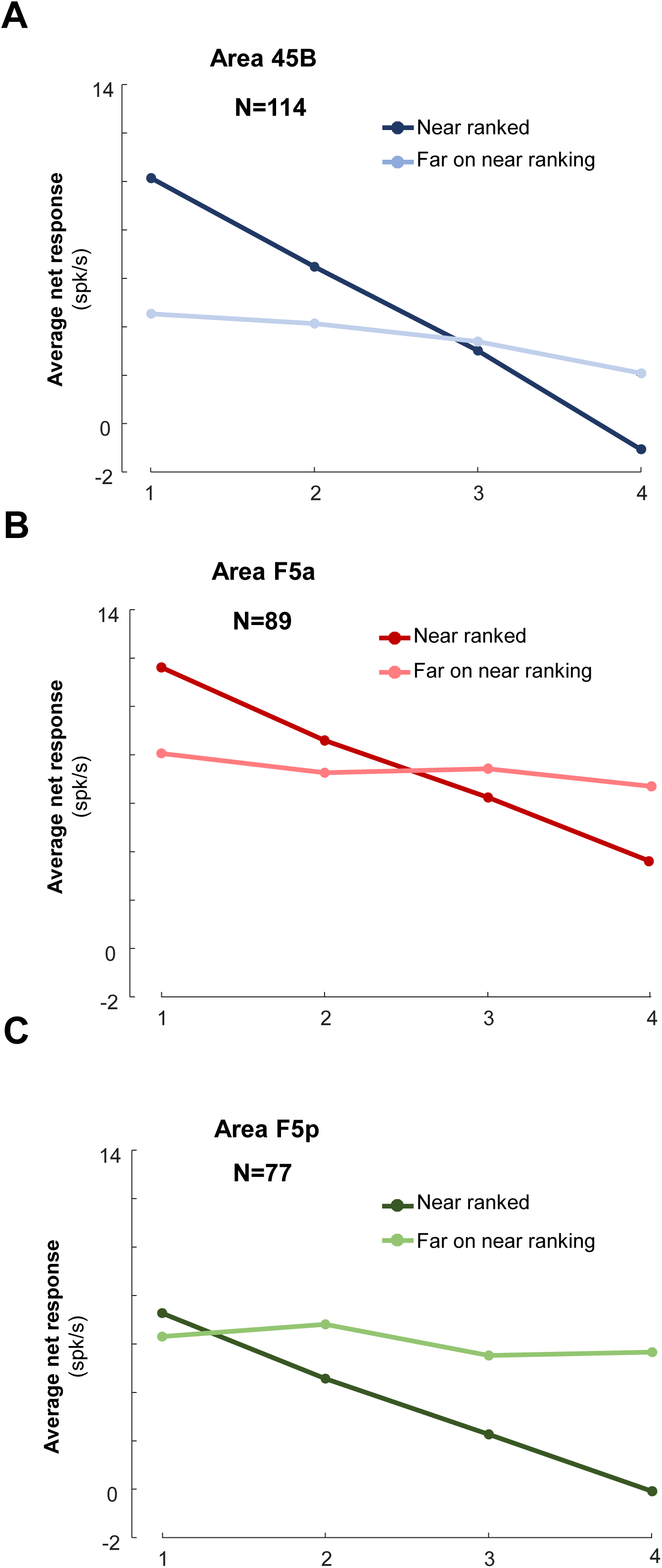
Ranking analysis: position tolerance across distances in depth. Average ranked responses at the Near position (dark color shades) and average responses to the same objects at the Far position (lighter color shades), for area 45B (blue; A), F5a (red; B) and F5p (green; C).

Because the viewing distance affected the object preference in all three areas, we determined the preferred object according to the highest response at the Near location, and compared this to the response to the same object at the Far position (Figure 7A), and vice versa (preferred object based on the Far location, Figure7B). Both during the early and the late epochs, area 45B and area F5a showed a strong effect of viewing distance (45B: early and late epochs, respectively p = 6.55 × 10^−5^ and 3.98 × 10^−7^; F5a: early and late epochs, p = 3.0 × 10^−3^ and p = 1.17x 10^−5^, respectively). Unexpectedly, however, area F5p had a weaker Near preference in the early epoch (p = 6.2 × 10^−3^), and no effect of viewing distance in the late stage of the trial (p > 0.05 – Figure 7A).When selecting the preferred object based on the Far location (Figure 7B), we observed a weak effect of viewing distance in 45B, which was much smaller than when selecting based on the Near responses (ANOVA, interaction effect between the Selected distance for ranking and preferred-nonpreferred distance p = 6.7 × 10^−3^). In F5p, however, the viewing distance effect was much stronger (ANOVA, p = 8.1 × 10 ^-3^), consistent with the higher proportion of Far preferring neurons. F5a showed an intermediate pattern of distance selectivity (moderate effect of viewing distance when selecting based on the Near responses, and a non-significant effect of viewing distance when selecting based on the Far responses, ANOVA interaction effect p = 0.80).

**Figure 7.**
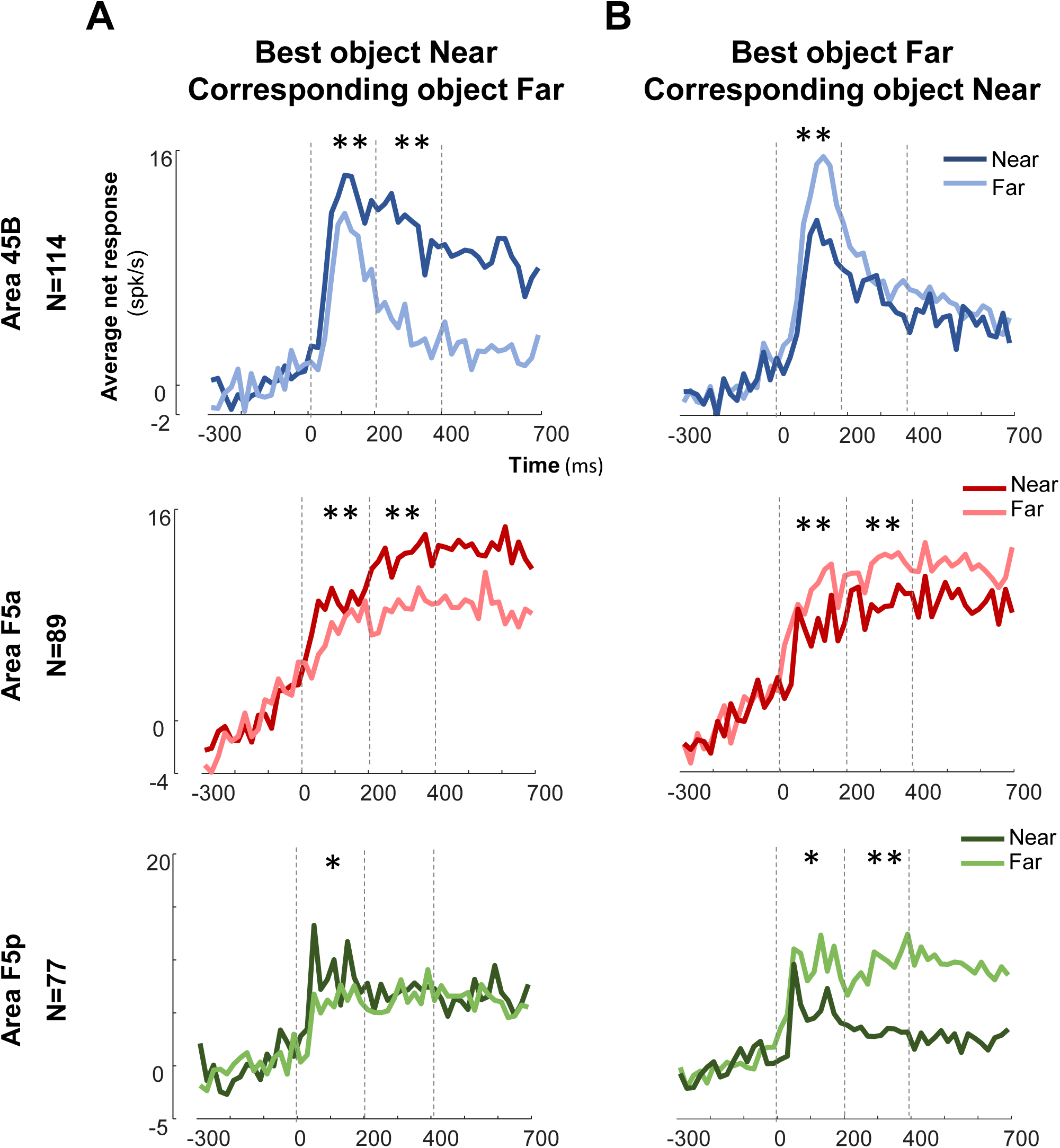
Distance effect with Near and Far neuron selection. A. Best object Near vs Corresponding object Far. B. Best object Far vs Corresponding object Near. For both A and B, bin size was 20 ms, the dark colors represent Near condition, and light color Far condition (blue, red and green, respectively for area 45B, F5a and F5p). Two asterisks one asterisk corresponds to a p<0.05; correspond to p<0.01.

Presenting the same object at the two positions also introduces a change in retinal size (Figure 2C), which may influence the neuronal responses. To investigate the effect of viewing distance for objects with identical retinal size, we compared the average net responses to the small object at the Near distance with those to the same shaped large object at the Far distance (i.e. small sphere Near vs large sphere Far, or small plate Near vs large plate Far – Figure 8). Only area 45B and F5a preserved a significant preference for the Near position during the Late epoch (t-test, p = 6.4 × 10^−3^ and p = 1.8 × 10^−3^, respectively), but not in the Early epoch (p = 0.23 for 45B and p = 0.07 for F5a). In contrast, F5p neurons did not distinguish between these two conditions in none of the epochs (p = 0.10 and p = 0.68 for Early and Late epoch, respectively). Therefore, for stimuli with identical retinal size, 45B and F5a neurons preserve their preference for the Near viewing distance, while F5p neurons show no clear preference anymore under these conditions.

**Figure 8.**
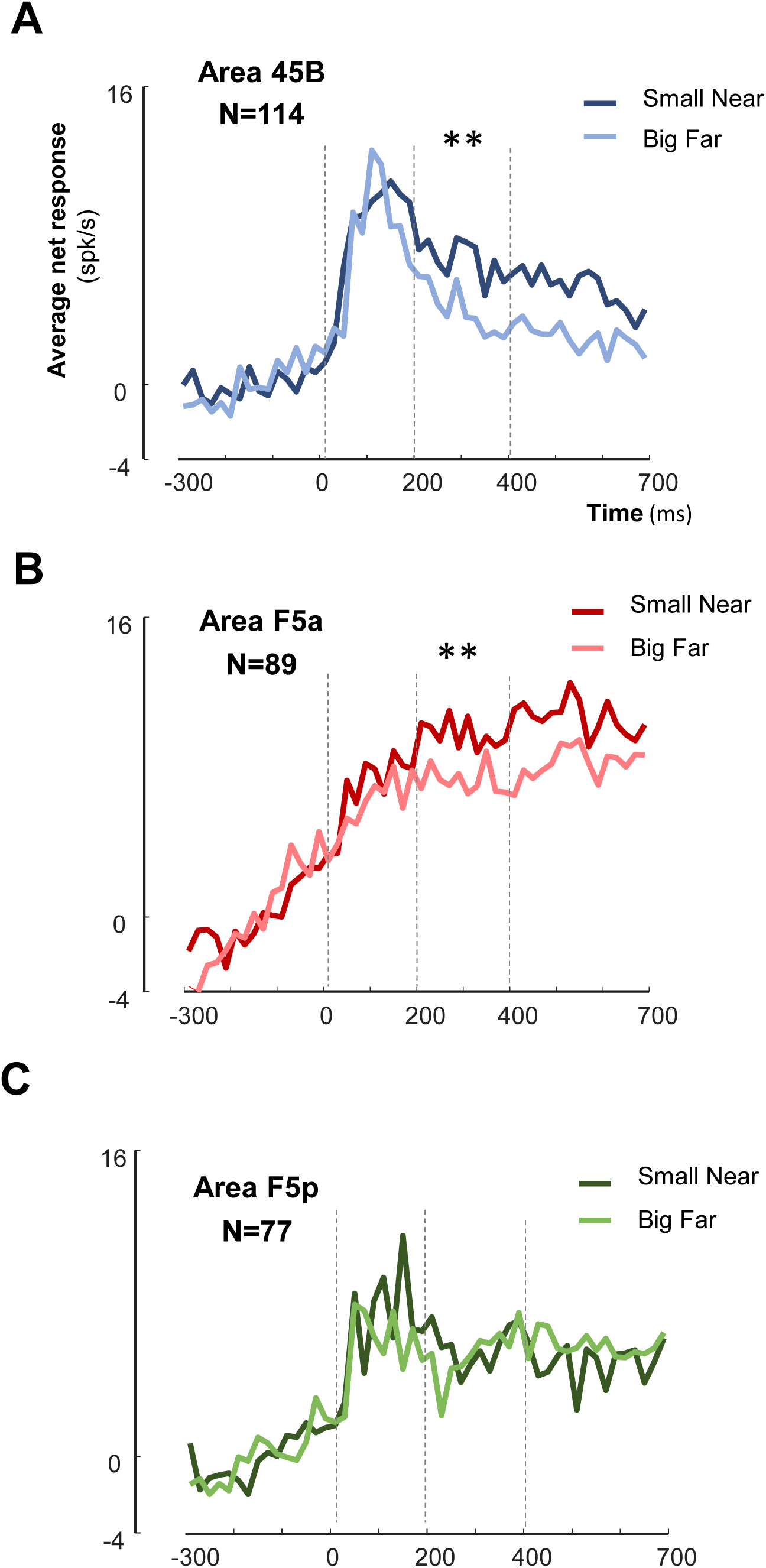
Distance selectivity controlled for retinal size. The objects compared have the same retinal size (i.e. Small Near object = Large Far object). Dark color represent small objects presented at Near position, while lighter color represent large objects presented at Far conditions (blue, red, and green respectively). Bin size = 20 ms. Two asterisks correspond to p<0.01.

### Grasping activity

We also recorded neural responses during visually guided grasping of the same objects. The majority of neurons that responded to Fix trials after Light onset, also responded in at least one epoch after Light onset in the grasping trials (Table 4). Figure 9B shows the average response to the best object aligned to Light onset, Go cue, Lift of the hand and Pull, respectively, in the VGG task (grasping in the light). As a general trend, we observed that area 45B (N = 114) and F5a neurons (N = 89) showed a response to Light onset, and a sustained response throughout the delay period and during the Pull of the object. In area F5p (N = 77), instead, we observed a transient response to Light onset, followed by a decrease in activity 200-300 ms after Light onset. Immediately after the Go-cue and before the Lift of the hand, the F5p activity rose strongly ([0-300] ms after lift of the hand compared to [300-0] ms before the Go cue, p = 4.03 × 10^−7^) and peaked around the Pull of the object. Thus, neurons in all three frontal areas are highly active during object grasping under visual guidance. As an additional test, we also recorded during a Memory guided grasping (MGG – Monkey Y – Figure 9A) task in a subset of the neurons (12 neurons in 45B, 5 in F5a and 6 in F5p – Figure 9C), in which the monkey had to reach and grasp an object in the dark. On average, neurons in 45B responded after Light onset, and became inactive in the delay period before the Go cue. In contrast, we observed sustained activity during MGG in F5a and F5p. Thus, the pattern of activity during VGG and MGG also indicates that the grasping activity in 45B is predominantly visual, whereas F5a and F5p also contain visuomotor and motor dominant neurons.

**Table 4.**
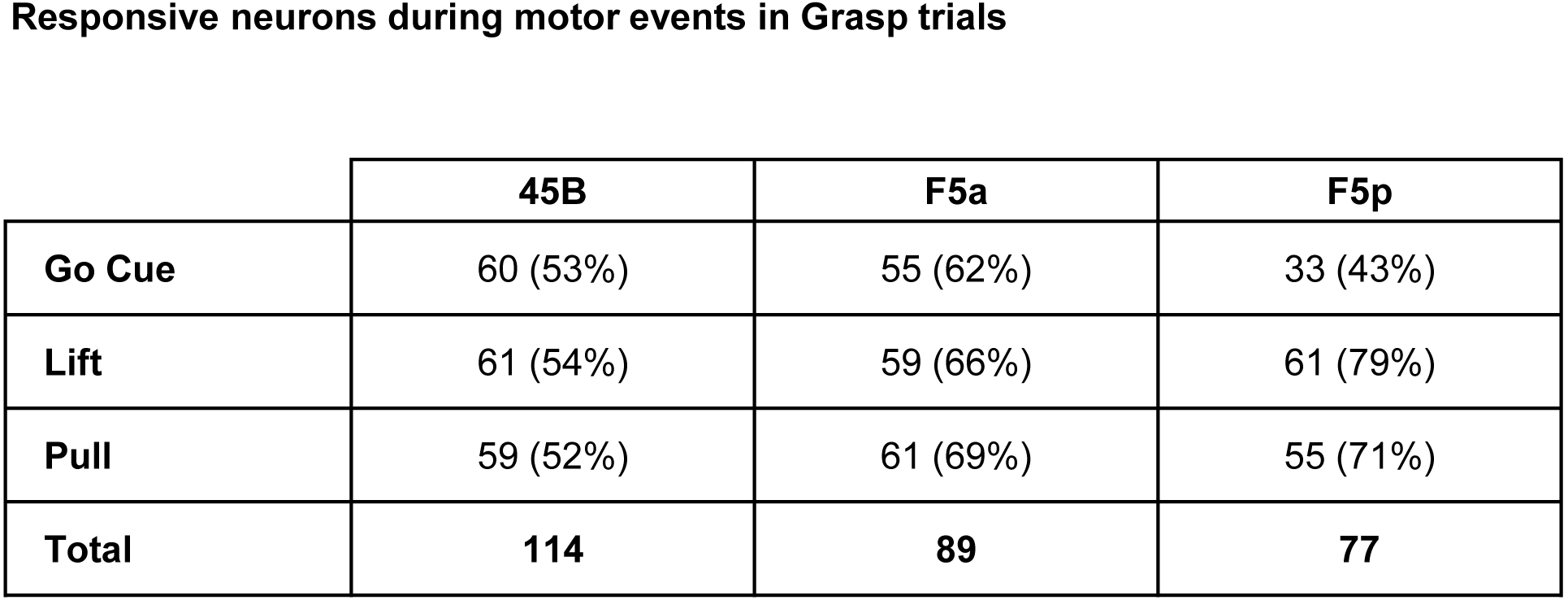
Percentage of responsive neurons during Go cue, Lift, and Pull events in Grasp trials

|  | **45B** | **F5a** | **F5p** |
| --- | --- | --- | --- |
| **Go Cue** | 60 (53%) | 55 (62%) | 33 (43%) |
| **Lift** | 61 (54%) | 59 (66%) | 61 (79%) |
| **Pull** | 59 (52%) | 61 (69%) | 55 (71%) |
| **Total** | **114** | **89** | **77** |

**Figure 9.**
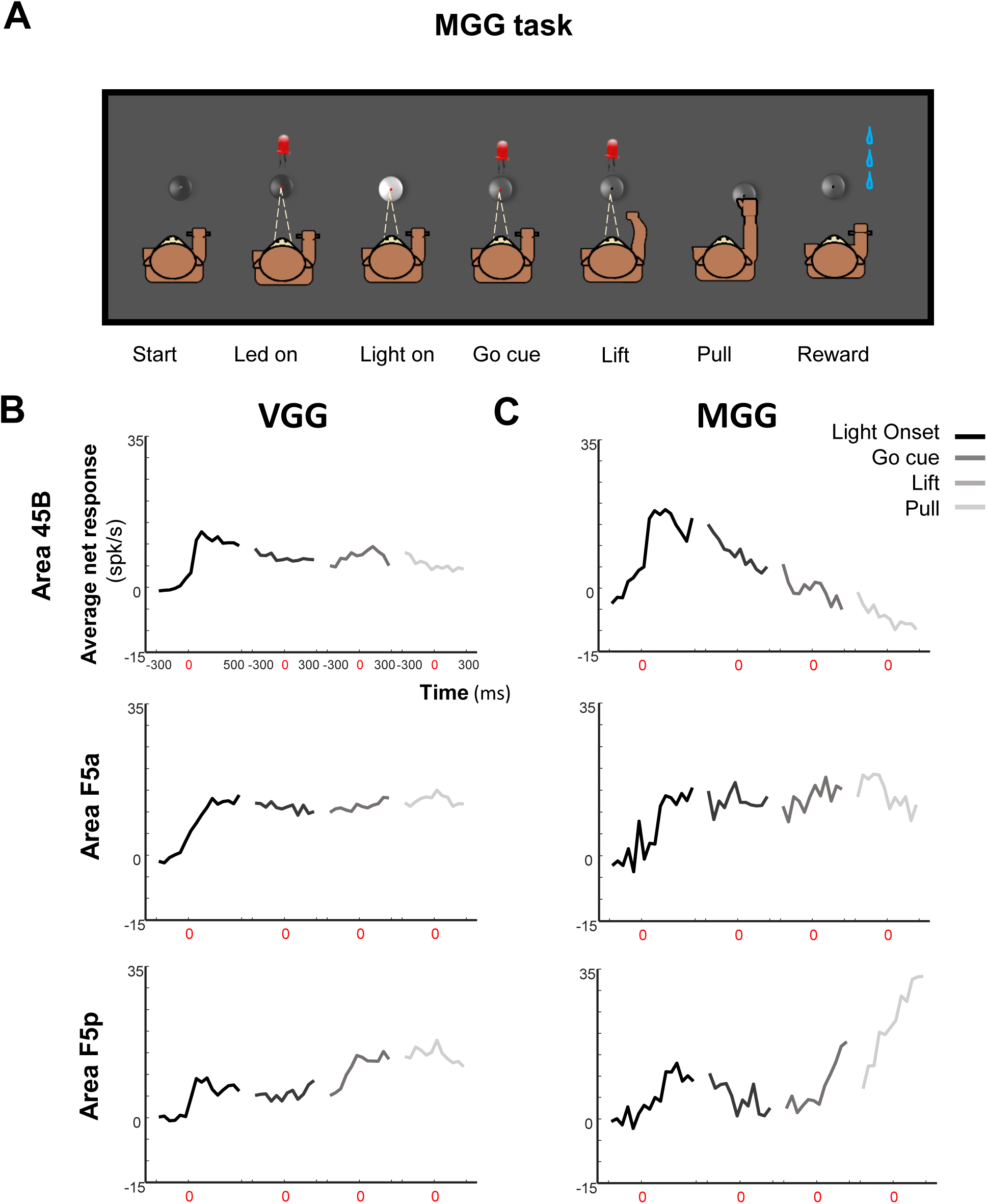
Grasping responses in area 45B, F5a and F5p. A. MGG tsk sequence. B. Average net grasping responses to the best object (Monkey D and Monkey Y), aligned to Light onset (black), Go cue, Lift and Pull of the object (progressively lighter color shades) for the three areas during the execution of the VGG task. Bin size = 50 ms. Spacing between alignments = 100 ms. Vertical dashed lines represent the multiple alignments. C. Average population responses in the three areas (Monkey Y) during the execution of the MGG task, respectively Area 45B (n =12), Area F5a (n = 5) and Area F5p (n = 6). Bin size 50 ms.

## 5. Discussion

We investigated the coding of object and viewing distance in area 45B, F5a and F5p using a task with interleaved passive fixation and VGG trials. Neurons in all three areas responded to the presentation of the object and were selective for viewing distance, but only 45B neurons were selective for both object and viewing distance. Contrary to our expectations, we observed a strong preference for the Near viewing distance in 45B and F5a and a preference for the Far viewing distance in F5p. Even for objects with identical retinal size at the two viewing distances, 45B and F5a neurons preferred the Near viewing distance.

The neural coding of viewing distance has been studied in early visual areas (Trotter et al., 1992), in dorsal (Andersen and Mountcastle, 1983; Gnadt and Mays, 1995; Genovesio and Ferraina, 2004) and in ventral stream areas (Dobbins et al., 1998). However, very few studies have investigated the neural coding of distance in the context of a reaching or grasping task. Ferraina et al. (2009) investigated the effect of binocular eye position in rostral parietal area PE on the reaching-related activity of individual neurons, and reported that a small subpopulation of neurons was influenced by viewing distance. Hadjidimitrakis et al. (2014, 2015) described joint selectivity for fixation distance and reach direction in the caudal parietal area V6A. In these studies, the fixation distance varied within the peripersonal space of the animal (i.e. less than 25 cm from the animal), and therefore no data were obtained for targets that appeared beyond reaching distance.

Similar to our previous study (Caprara et al., 2018), area 45B neurons showed fast and selective visual responses to the object after Light onset. Previous studies described fMRI activations in this area evoked by 2D images of objects (Denys et al., 2004; Nelissen et al., 2005), and selectivity of individual neurons for shapes and very small line fragments (Caprara et al., 2018), and zero order disparity (Theys et al., 2012). With the current results, we confirmed the involvement of these neurons in shape and object processing. Most 45B neurons were also significantly affected by viewing distance and showed a preference for Near, even when correcting for retinal size.

Because of the visual properties of this area and the direct anatomical connection with pAIP (Premereur et al., 2015), we previously hypothesized that area 45B could be involved in oculomotor control, similar to the neighboring region FEF (Gerbella et al., 2010; Caprara et al., 2018), to guide eye movements towards specific parts of the object contour. However, we observed sustained object responses in a visually-guided grasping task, which does not seem to be consistent with pure oculomotor control since no saccade was required after obtaining fixation. Our results rather suggest that area 45B may have a role in object processing and eye-hand coordination when grasping objects under visual guidance. Consistent with this interpretation, the activity during grasping in the dark was very low in 45B. In addition to this, the strong distance selectivity with a preference for Near, was also inconsistent with our initial hypothesis, and could not be explained as a vergence effect, as the eye position was stable after object onset (data not shown). At least for the subpopulation of neurons preferring the peripersonal space, we cannot exclude the possibility that the distance effect we observed was related to the significance that the stimulus acquired when it was reachable, and therefore graspable. Our results are in accordance with those of the fMRI study by Clery et al. (2018), who reported activations in prefrontal cortex in response to object presentation at reachable distances.

The results we obtained in F5a were largely consistent with the known anatomical connectivity and neuronal properties of this area. Although we measured significant object selectivity in individual F5a neurons, our population did not discriminate reliably between the four objects, most likely because we only used a limited number of objects. Moreover, the object responses we measured were relatively slow and were preceded by clear anticipatory activity in the epoch immediately before light onset. Nevertheless, the F5a neurons we recorded were strongly affecting by viewing distance, preferred the Near viewing distance even for stimuli with identical retinal size, and were highly active during visually-guided and during memory-guided object grasping (a mixture of visual-dominant and visuomotor neurons as demonstrated in Theys et al. 2013). Overall, these properties are consistent with the proposed position in the cortical hierarchy as ‘pre-premotor cortex’ (Gerbella et al., 2010), receiving visual inputs from AIP and transmitting information to F5p.

Previous studies have described the connectivity pattern of F5p as strongly motor-oriented, receiving input from F5a and projecting directly to primary motor cortex (Borra et al., 2010; Gerbella et al., 2011). Being part of the same pathway (pAIP ← aAIP ← F5a ← F5p), several authors compared object representations in AIP and F5p (Murata et al., 2000; Raos et al., 2006). They concluded that AIP provides a visual object description, while area F5p represents the same object in motor terms, i.e. the grip type necessary to grasp the object Fogassi et al. (2001) confirmed the strong motor character of this area by reversibly inactivating F5 (probably F5p) with muscimol, which induced a deficit in the preshaping of the hand during grasping. Our population of neurons showed a clear involvement in grasping in the light and in the dark (Figure 9A). Because objects may be encoded in motor terms in F5p, we hypothesized that F5p would strongly prefer the Near distance but unexpectedly, a high number of neurons preferred the Far viewing distance. At first glance, our results seem to be in contrast to those of Clery et al. (2018), reporting Near preference in F4/F5p area. However, the ‘Far space’ distance for object presentation used by (Clery et al., 2018) was significantly larger than in our case (150 cm and 56 cm, respectively). Therefore, we believe it could be possible that our Far viewing distance was not long enough to reduce the visuomotor neural responses in F5p. Another difference with our study is that Clery et al. (2018) analyzed fMRI responses in monkeys, which may also include subthreshold modulations and presynaptic activity (Logothetis, 2008).

Our results are important for the interpretation of the organization of the parieto-frontal grasping network. At the level of the posterior subsector of AIP, visual information is transmitted along two parallel pathways: towards anterior AIP, and then to F5a and F5p, and directly to 45B (Premereur et al., 2015). Importantly, the strong object – and distance selectivity we observed in 45B, together with its activity during VGG but not during MGG, could be observed in any visual area in occipital, temporal or prefrontal cortex, but does not by itself indicate a causal role in object grasping. If 45B would be found to be causally related to object grasping, we believe that eye-hand coordination (monitoring the position of the own hand approaching the object) may be the most likely aspect of object grasping supported by 45B neurons.

In conclusion, our data suggest a much more complex role of 45B in the network rather than directing saccades towards object contours. Moreover, the strong visual responses and the surprising preference for the Far viewing distance of area F5p suggest that the visuomotor object representation in F5p was still activated when we presented objects beyond reaching distance. Although not tested in the current experiment, the presence of a transparent barrier interposed between the monkey and the object could have decreased or silenced the F5p response to objects located both at the Near and Far distances in a similar way as in Bonini et al. (2014). Further experiments will have to investigate the causal role of these areas (specifically area 45B and F5a) in the grasping network. We expect that pharmacological approaches such as muscimol reversible inactivation will be able to shed light on the causal role of these three areas in visuomotor transformations during object grasping.

## Acknowledgements

This work was supported by Fonds voor Wetenschappelijk Onderzoek Vlaanderen (Odysseus grant G.0007.12 and grant G0A8516N). We thank Stijn Verstraeten, Piet Kayenbergh, Gerrit Meulemans, Marc De Paep, Astrid Hermans, Christophe Ulens and Inez Puttemans for assistance, and Steve Raiguel for comments on a previous version of this manuscript.

